# Density-by-diet interactions during larval development shape adult life-history trait expression and fitness in a polyphagous fly

**DOI:** 10.1101/2021.11.29.470426

**Authors:** Juliano Morimoto, Anh The Than, Binh Nguyen, Ida Lundbäck, Hue Dinh, Fleur Ponton

## Abstract

Habitat quality early in life determines individual fitness, with possible long-term evolutionary effects on groups and populations. In holometabolous insects, larval ecology plays a major role in determining the expression of traits in adulthood, but how ecological conditions during larval stage interact to shape adult life-history and fitness, particularly in non-model organisms, remains subject to scrutiny. Consequently, our knowledge of the interactive effects of ecological factors on insect development is limited. Here, using the polyphagous fly *Bactrocera tryoni*, we conducted a fully-factorial design where we manipulated larval density and larval diet (protein-rich, standard, and sugar-rich) to gain insights into how these ecological factors interact to modulate adult fitness. As expected, a protein-rich diet resulted in faster larval development, heavier and leaner adults that were more fecund compared with standard and sugar-rich diets, irrespective of larval density. Females from the protein-rich larval diet had overall higher reproductive rate (i.e., eggs per day) than females from other diets, and reproductive rate decreased linearly with density for females from the protein-rich but non-linearly for females from the standard and sugar-rich diets over time. Surprisingly, adult lipid reserve increased with larval density for adults from the sugar-rich diet (as opposed to decreasing, as in other diets), possibly due to a stress-response to an extremely adverse condition during development (i.e., high intraspecific competition and poor nutrition). Together, our results provide insights into how ecological factors early in life interact and shape the fate of individuals through life-stages in holometabolous insects.

## Introduction

Habitat quality early in life (i.e., ‘developmental environment quality’) can determine the fitness of individuals in the current (direct) and next generations (indirect), thereby regulating physiological and behavioural responses as well as population dynamics and evolution (Mousseau and Fox 1998; Wolf et al. 1998). Studies in both invertebrate and vertebrate species have shown that the quality of the developmental environment can influence fitness traits in adulthood [e.g., (Booth and Wellington 1998; Stevens et al. 1999, 2000; Steigenga and Fischer 2009; Spencer et al. 2010; May et al. 2015; Stefana et al. 2017; Eyck et al. 2020)], canalising individuals into strategies that in turn affect population dynamics (Booth 1995; Lindström 1999; Gaillard et al. 2000; Monaghan 2008). Herbivorous holometabolous insects are ideal to study individual- and population-level effects of developmental ecological factors. This is because ecological factors that influence the quality of larval developmental environment in herbivorous insects are relatively simple to model [e.g., (Atkinson and Shorrocks 1981)] and manipulate [e.g., (Morimoto et al. 2018)], and display long-lasting quantifiable effects on fitness [e.g., (Rossiter 1992; Steigenga and Fischer 2009; Takken et al. 2013; Tigreros 2013; Matavelli et al. 2015; Rodrigues et al. 2015; Morimoto et al. 2016, 2019b; Schwab et al. 2017; Than et al. 2020)]. This is ideal for controlled experimental approaches that envisage to test theories and provide new empirical evidence to guide further ecological research assessing the quality of developmental environments, particularly because the concept and scope of (developmental) habitat can be complex to define and properly measure in uncontrolled environments (Mitchell 2005). In fact, two ecological factors are known to be major modulators of the quality of the larval developmental habitat of herbivorous holometabolous insects: conspecific density (i.e., intraspecific competition) and larval diet composition (Jaenike 1978; Than et al. 2020). While conspecific larval density can (but not always does) increase protection against predators (Rohlfs and Hoffmeister 2004), it also increases intra-specific competition thereby decreasing resource availability (Atkinson and Shorrocks 1984; Klepsatel et al. 2018), resulting in an inverse relationship with adult reproduction and population growth (i.e., ‘density-dependent effects’) [e.g., (Capinera and Barbosa 1977; Gage 1995; Baldal et al. 2005; McGraw et al. 2007; Morimoto et al. 2016, 2017; Wigby et al. 2016)]. Likewise, larval nutrition modulates larval growth as well as adult size, reproduction, and the expression of other fitness traits (‘diet-dependent effects’) (Cookman et al. 1984; Greene 1989; Perkins et al. 2004; Matavelli et al. 2015; Rodrigues et al. 2015; Silva-Soares et al. 2017; Jang and Lee 2018; Morimoto et al. 2019b, 2020; Shrader et al. 2019). Therefore, herbivorous holometabolous insects can be excellent models to test the long-term consequences of the developmental environment in controlled experiments, from which the results can help advance ecological theory and thus be generalised to broader ecological contexts.

While previous studies have addressed the role of either larval diet or density individually, few have investigated the interactive effect of these ecological factors on insect fitness. Nonetheless, the few studies that did investigate the interaction are overall consistent in their effect: (i) increasing larval densities increase developmental time while decreasing adult body size and fecundity whereas (ii) nutrient-rich diets, particularly rich in protein, have the opposite effect. This is because larval crowding is thought to deplete protein availability in the diet, thereby becoming a limiting growth factor for larval development; a diet that is protein-rich can in theory counterbalance this effect, and allows for larval growth even in crowded conditions (Klepsatel et al. 2018). For instance, black soldier fly larvae *Hermetia illucens* are particularly lighter and have relatively lower larval fat reserves in nutrient-poor and crowded developmental environments (Barragan-Fonseca et al. 2018). Similar effects have been observed in *Aedes aegypti*, whereby the negative relationship between larval developmental time and larval crowding is steeper in nutrient poor diets (inferred from the test statistics, as no display of the significant interaction was given) (Couret et al. 2014). In the Queensland fruit fly (*Bactrocera tryoni*), sugar-rich semi-sterile diets have a disproportionate negative effect on pupal weight in crowded environments (Nguyen et al. 2019) while in *Drosophila melanogaster*, increasing yeast content in the diet of crowded larvae rescues adult survival and fat reserves (Klepsatel et al. 2018). Thus, overall, there has been empirical data showing that larval density and diet interact, and that diet can rescue life-history trait expressions caused by crowding. However, many questions remain. Firstly, although empirical data suggest that crowding and nutrient content (particularly protein content) have single opposite effects on fitness, the above studies also found important but unexplored interactions between crowding and diet. For instance, in *H. illucens*, the decrease in larval weight with increasing larval density is more pronounced in nutrient-poor diets compared with nutrient-rich diets (Barragan-Fonseca et al. 2018). Furthermore, while nutrient content may rescue physiological traits and lifespan of individuals from high-density developmental environments (Klepsatel et al. 2018), it is still unknown if the paths through which individuals attain such fitness are similar. For instance, small *D. melanogaster* populations composed of individuals from crowded larval environments invest in reproduction early in life, whereas groups composed of individuals from uncrowded larval environments display a maximum reproduction at almost the half-life of the group (Morimoto et al. 2017). Yet, lifetime fitness (*r*) of groups differs only slightly, suggesting that individuals and groups can attain similar fitness levels through different physiological responses. Whether these differences emerge independently from signals triggered by direct intraspecific competition levels (i.e., crowding), indirect nutritional signals (e.g., protein depletion), or the interaction between both factors remains to be investigated. More broadly, it is still unclear how plastic responses to the interactions between diet and intraspecific competition level modulate life-histories and fitness of individuals. Individuals and populations are likely to experience fluctuating developmental conditions over time as well as over their distribution range [see e.g., (Singer and McBride 2012)] for which plastic responses can be a key factor in supporting population survival in novel conditions (Hendry, 2016). Therefore, studying how the synergistic interactions between diet composition and intraspecific competition levels interact to shape individuals’ life-histories and fitness is crucial for understanding long-term eco-evolutionary processes such as adaptation to novel environments and diet specialisation.

To address this gap, we used the herbivorous fruit fly *Bactrocera tryoni* (Diptera: Tephritidae) as model to investigate the effects of conspecific density and nutrition on life-history trait expression and fitness. We performed a controlled fully-factorial experiment in which we created five larval densities ranging from very low to very high across the three diets (i.e., protein-rich, standard (balanced) and sugar-rich diets), measuring the interactions between diet- and density-dependent effects on developmental time, pupal and adult weights, adult energetic reserves and fecundity. These traits have been shown to respond to larval density and/or crowding (Than et al. 2020), and therefore are suitable proxies to assess our experimental manipulation effects on fitness-related traits. *Bactrocera tryoni* has an interesting natural history which makes this a good system for our study. For instance, females *B. tryoni* use a wide variety of hosts species to oviposit (Hancock et al. 2000). Eggs are deposited in small batches of 4-20 eggs per oviposition bout, and several females lay eggs in the same fruit so that offspring densities can vary (Fitt 1986, 1990). In addition, larvae are known to aggregate in a diet-dependent fashion with potential fitness benefits but also with costs of increasing intraspecific competition (Christenson and Foote 1960; Morimoto et al. 2018). Based on ecological theory and previous studies of density- and diet-dependent effects, we obtained the predictions highlighted in Table 1. Our study addresses a current but unexplored gap in our knowledge about how nutrition and population density can interact to shape fitness, using a polyphagous fly of ecological and economic significance as model. The findings presented here highlight the need for a more integrative view of developmental ecology of (holometabolous) insects.

**Table 1.** Predictions.

|  | **Prediction** | **Literature** |
| --- | --- | --- |
| 1 | Increasing densities would decrease fitness-trait expression (e.g., fecundity) due to increased intraspecific competition | (Courchamp et al. 1999) |
| 2 | Protein-rich diets would result in higher average fitness-trait expression (e.g., higher adult fecundity) compared with sugar-rich and, to a lesser extent, standard (balanced) diet | (Matavelli et al. 2015; Silva-Soares et al. 2017; Nguyen et al. 2019) |
| 3 | Sugar-rich diets would result in higher lipid storage in adulthood, while protein-rich diets would generate individuals with the lowest lipid reserves | (Musselman et al. 2011; Morimoto et al. 2020) |
| 4 | If larval density effects are generated solely by dietary protein limitation, there should be an interaction between diet and density highlighting the overall higher performance of individuals across densities in protein-rich diets | (Klepsatel et al. 2018) |

## Materials and methods

### Fly stock

Eggs were collected from females in our laboratory-adapted stock of *B. tryoni* that was established with flies collected in Ourimbah, NSW in 2015 (>30 generations old). Our stock has been maintained in non-overlapping generations. For the stock, larvae were given gel-based diet as in Moadeli, Taylor and Ponton, (2017) and adults were provided a free-choice diet of hydrolysed yeast (MP Biomedicals Cat. n° 02103304) and commercial refined sucrose (CSR^®^ White Sugar). We collected eggs for 1h in a 300 mL semi-transparent white soft plastic bottle (low density polyethylene) that had numerous perforations of <1mm diameter through which females could oviposit. To maintain humidity, the bottle contained 20 mL of water. All stocks and experiments were maintained in a controlled environment room at 65 ± 5% relative humidity and 25 ± 0.5°C, with light cycles of 12 h light: 0.5 h dusk: 11 h dark: 0.5 h dawn in the Department of Biological Sciences at Macquarie University.

### Diets

Three diets were used for all experiments which varied in yeast-to-sugar ratios (i.e., Y:S ratio). The details of the diets have been described in detail in Nguyen *et al*., (2019). Briefly, the standard diet followed the original recipe [see (Moadeli et al. 2017)] and had Y:S ratio of 1.6:1, the protein-rich diets had Y:S ratio of 4:1 and the sugar-rich diet, Y:S ratio of 1:3 (Nguyen et al. 2019). For preservation of the diets, we also included Nipagin (Southern Biological^®^ cat no. MC11.2), Sodium Benzoate (Sigma^®^ cat no. 18106), and Critic Acid (Sigma^®^ cat no. C0759) for all experimental diets. All ingredients were boiled in a 1% water-agar solution and poured into small (55 mm x 16 mm) and large (100 mm x 20 mm) Petri dishes which were allowed to set at room temperature for 10 minutes before being closed and individually wrapped into cling film for storage at 4°C prior to the experiments.

### Experimental design

Eggs were collected for two hours from the stock colony as described above. Based on our previous protocol to manipulate larval density [e.g.,(Nguyen et al. 2019)], we deposited different volumes of egg-water solution onto the diets to generate five densities (D) of larvae: <u>D1</u>: 8 μL of egg solution (~110 larvae/mL diet); <u>D2</u>: 16 μL of egg solution (~220 larvae/mL diet); <u>D3</u>: 30 μL of egg solution (~450 larvae/mL diet); <u>D4</u>: 65 μL of egg solution (~930 larvae/mL diet) and <u>D5</u>: 125 μL of egg solution (~1800 larvae/mL diet). These densities include the range of potential densities that Tephritidae larvae of many species (including *B. tryoni*) can encounter [e.g., (Fitt 1986)] and also extends larval densities values in both the low and high-density tails. We had seven replicates *per* diet *per* density (*N_total_* =105). After 5-6 days that the eggs were deposited in the diet, we open the lids and placed the Petri dishes into a 1.125L plastic cages with 30g of vermiculite for pupation. Pupae was sieved as described above, placed into a large Petri dish (100 mm x 20 mm) and separated to different performance experiments. Ten randomly selected pupae per replicate per diet per density (*N* = 1050) were weighed in a Sartorius^®^ ME5 scale (0.0001 g precision) and returned to the pool of pupae. Thirty randomly selected pupae per diet per density were placed into small Petri dishes (55 mm x 16 mm) and into 1.125L cages with water. We measured the time from the first larval jump to vermiculite to the first adult emergence (in days) as proxy for developmental time. This is an appropriate proxy because *Bactrocera tryoni* utilises ephemeral patches and display tightly regulated developmental time [e.g., (Kumaran and Clarke 2014; Morimoto et al. 2019b)], as seen in other taxa that explores ephemeral patches during development [e.g., (Morey and Reznick 2004)]. Percentage of adult emergence was estimated as the number of fully emerged adults divided by the number of pupae (i.e., 30) x 100. Adults were collected and stored in −20°C within five hours post-emergence for body weight and lipid percentage measurements. This was done as following: five randomly selected males and five randomly selected females per replicate per diet per density (*N* = 1050) were placed individually in 10 mL glass tubes and dried at 60°C for three days in a drying oven, after which dried bodies were weighed on a Sartorius^®^ ME5 scale. Two mL of chloroform (Sigma Aldrich^®^, Cat no. 288306) was then added to each tube which was then sealed with a rubber plug and held for 24 h before the chloroform was discarded; this process was repeated once a day for three consecutive days. Bodies were then placed in the fume hood for 48h after which they were dried again at 60°C for three days in a drying oven before we measured body weight after lipid extraction [as in (Morimoto et al. 2019a; Nguyen et al. 2019)]. The percentage of body lipid was calculated as the difference between the dry body weight before and after lipid extraction, standardized by the body weight of each fly before the lipid extraction multiplied by 100 (i.e., percentage of lipid relative to the original dry body weight of each fly). Lastly, we also measured adult daily and total female fecundity as following: five recently-emerged males and five recently-emerged females were placed into a 1.125L plastic cage with *ad libitum* hydrolysed yeast, sucrose, and water. After 10 days post-emergence, we introduced the oviposition device which consisted of a small Petri dish (55 mm x 16 mm) covered with parafilm containing ~1 mm perforations and with 5mL of commercial apple juice to incite egg laying. Oviposition device was replaced daily for 20 consecutive days when females were 12 days-old (no oviposition was observed before this time). *Bactrocera tryoni* is anautogenous (Fisher 1994) and reach sexual maturation 10–12 days after emergence under laboratory conditions, meaning that our protocol allowed us to collect eggs throughout females’ peak reproductive performance and maximum contribution to population growth. For convention and clarity, we assigned day 0 to the day when females started laying eggs (sexual maturity) as this is the day when individuals contribute to population growth. We divided the number of eggs by the number of females in the group to have a standardised measure of ‘daily per female fecundity’. Total fecundity per female fecundity was then calculated as the sum of the daily per female fecundity per replicate per density per cage (*N* = 1050). A schematic representation of the experimental design is shown Fig 1.

**Figure 1.**
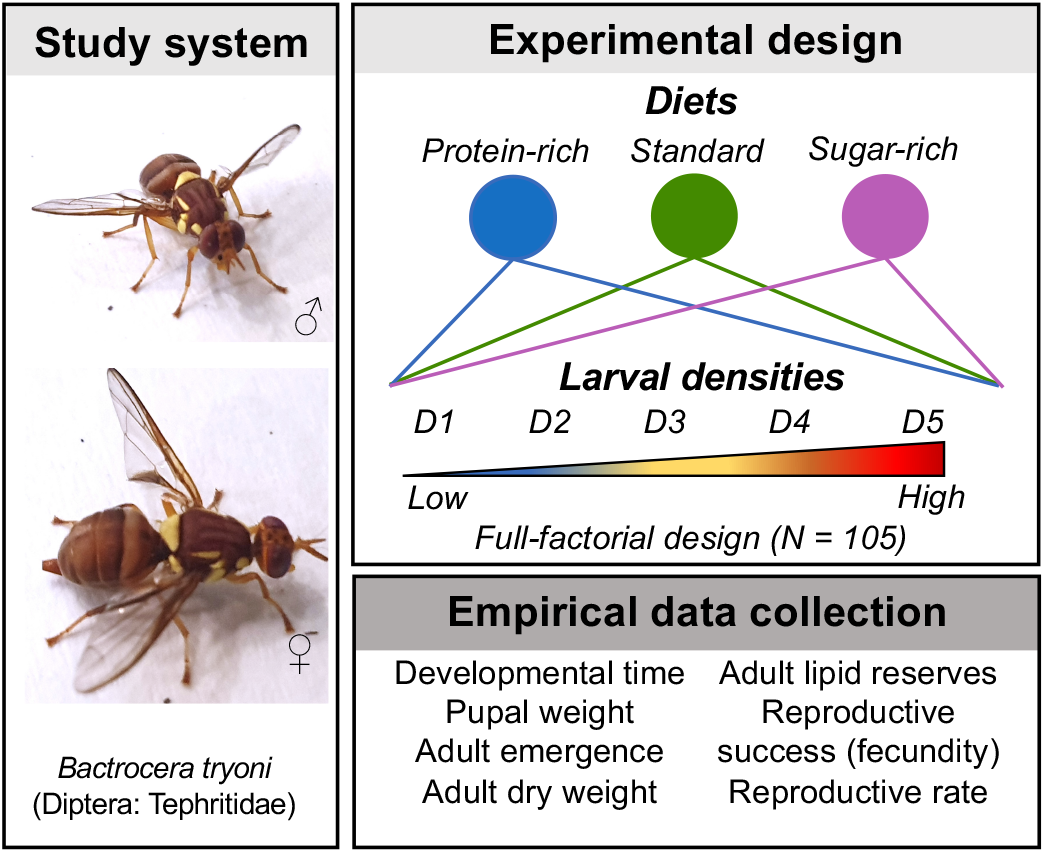
Experimental design and empirical traits measured. **(a)** Schematic representation of the fully-factorial experimental design whereby we manipulated diet composition (i.e., protein-rich, standard (balanced), and sugar-rich diets) and density (D1 = low to D5 = high) of *B. tryoni* larvae. We measured the life-history effects of larval diet and density during development. D1: ~110 larvae/mL diet; D2: ~220 larvae/mL diet; D3: ~450 larvae/mL diet; D4: ~930 larvae/mL diet and D5: ~1800 larvae/mL diet (see Methods for details).

### Statistical analyses

We used linear mixed models (LMM) from the ‘lme4’ and ‘lmerTest’ packages for statistical modelling and inferences (Bates et al. 2007; Kuznetsova et al. 2017), adding population ID as a random intercept in the model. Models for developmental time, pupal and adult weights, adult emergence, sex ratio, lipid reserves and reproductive success (i.e., total egg production) included diet, the linear and non-linear effects of density, and their interaction. When modelling adult traits, we also included sex and the two- and three-way interactions with diet and (linear and quadratic) densities. We also used LMM to investigate daily egg production per female (‘reproductive rate’), which included a random slope of population ID over time and the fixed effect interactions between diet and the linear effects of time and density as well as diet and the quadratic effects of time and/or density. For all models, we used the contrast comparison approach with the ‘standard’ diet as the reference level. In all models, we fitted density as a continuous variable even though we had five discrete treatments because this approach allowed us to study linear and non-linear effects of density on the dependent variables. We scaled density and day to improve fit of the models. For simplicity of presentation, we used the discrete densities D1, D2, D3, D4 and D5 as landmarks of lowest, low, intermediate, high, and highest densities for the presentation of the results and the discussion of our findings. It is important to mention that larval developmental time was highly synchronous in our controlled experiment, and thus variability was substantially reduced or absent in some treatment groups. For consistency, we nevertheless analysed the data with the mixed models approach described above, but p-values should be interpreted with caution for our model of developmental time. For the analysis of reproductive success, we removed an outlier with Cook’s distance > 0.5. While this removal qualitatively changed the significance of the interaction between diet and the non-linear effect of density (from F_2,95_ = 2.059, p = 0.133 to p < 0.047, see Table 2), this change does not qualitatively affect the conclusions of the study. We used AIC/BIC criterion to evaluate and select all models used for the analyses. All analyses were performed in R version 3.6.2 (R Core Team 2019). Scatterplots and boxplots were done using the ‘ggplot2’ package (Wickham 2016) while 3D surface plots were done using the ‘Tps’ function of the ‘fields’ package (Nychka et al. 2017).

**Table 2.**
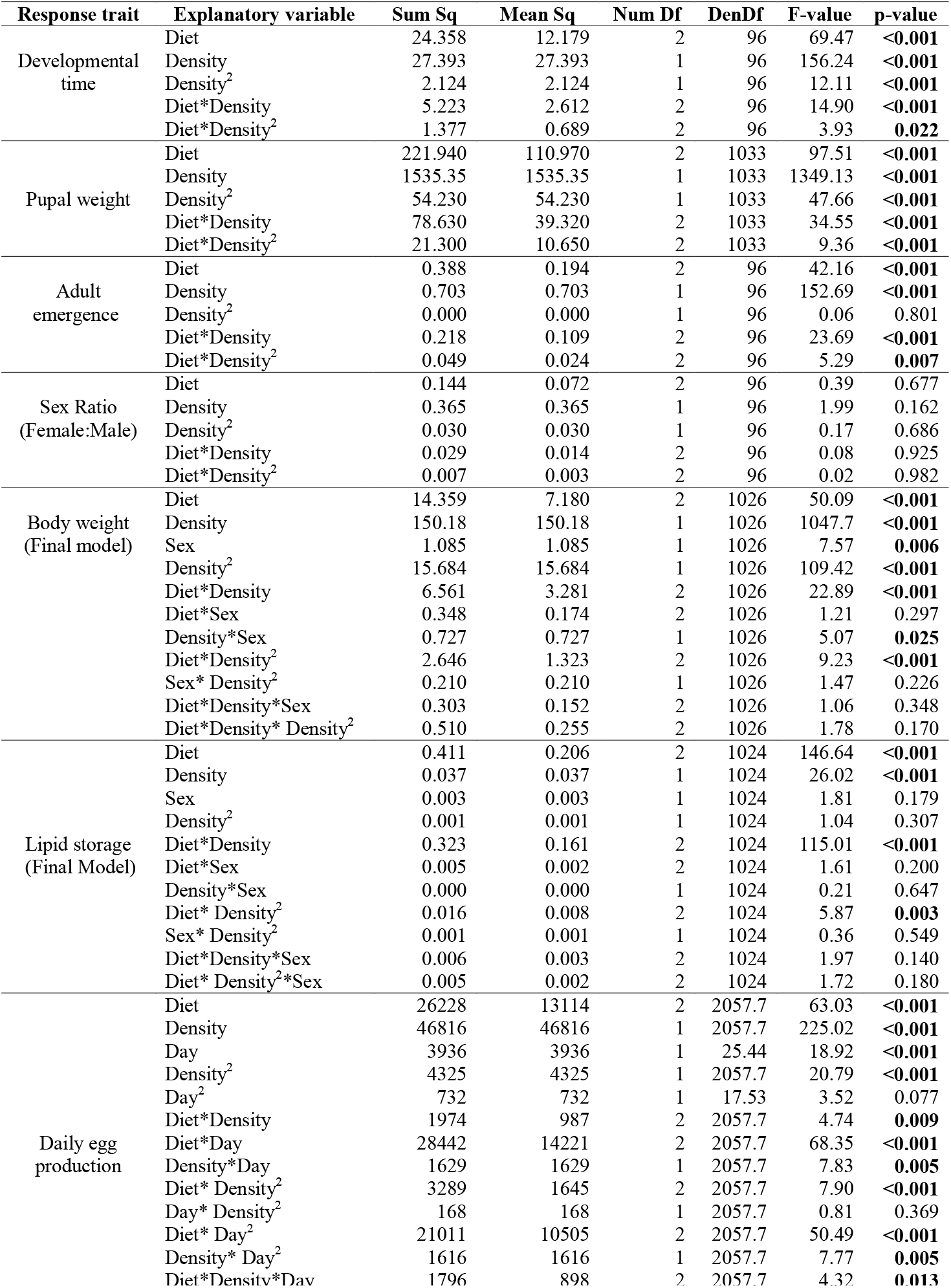
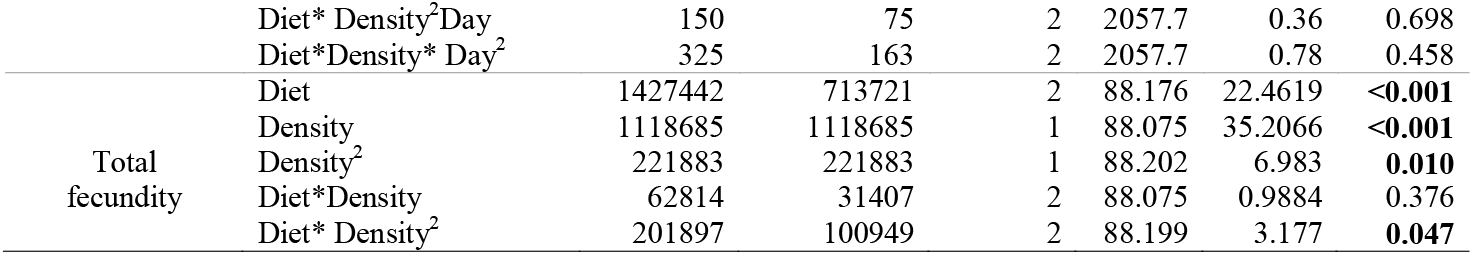
Complete LMM output of the models used for statistical inference in this study. Note that for these models, density was used as numeric, which allowed us to fit linear and non-linear effects (see Methods). **Bold:** p < 0.05.

## Results

### Density-by-diet effects on developmental traits

#### Developmental time

Relative to the standard and sugar-rich diets, larvae across all densities raised in protein-rich diets developed faster (*Diet*: F_2,96_ = 69.465, p < 0.001, Table 2, Fig 2a). Moreover, larvae in higher densities also developed faster than larvae in lower densities, and display an inflexion point (around D3) from which development was accelerated (*Density*: F_2,96_ = 156.24, p < 0.001; *Density^2^*: F_2,96_ = 12.113, p < 0.001). Importantly though, there were statistically significant interactions between diet and the linear and quadratic effects of density on developmental time (*Diet*Density^2^*: F_2,96_ = 3.921, p = 0.022; Table 2). These effects emerged because developmental time was shorter at high densities but this effect was more pronounced (stronger negative linear slope) in the standard diet compared to protein- and sugar-rich diets (Fig 2a). Moreover, while the non-linear inflexion point was around D3 for both standard and protein-rich diets, the inflexion was significantly less pronounced in the sugar-rich diet, where the relationship was close to linear (Fig 2a).

**Figure 2.**
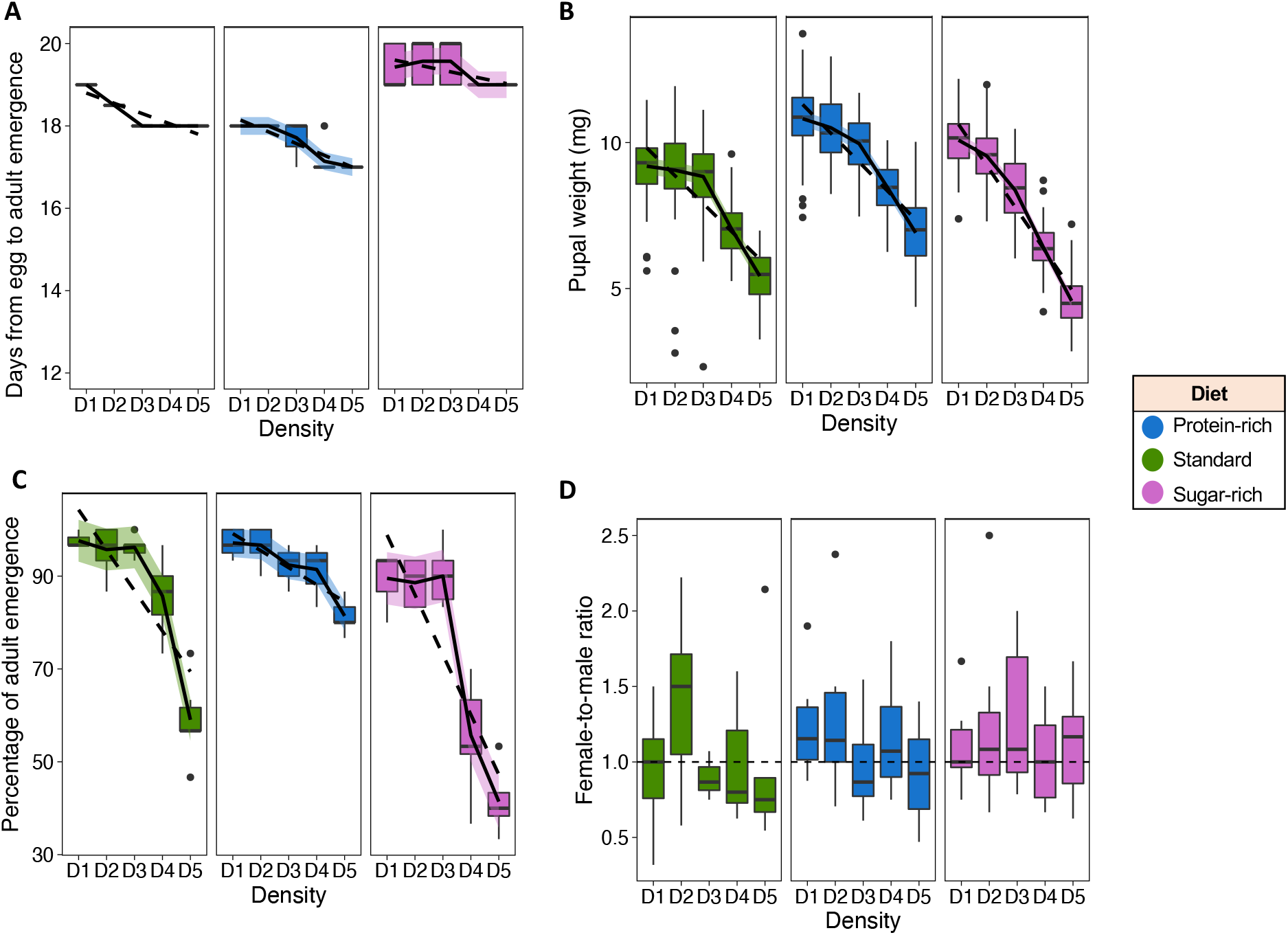
Density-by-diet effects on developmental traits. **(a)** Developmental time (egg-to-adult) in days. Note that for standard diet and to a smaller extent, protein- and sugar-rich diets, developmental time was highly synchronised. **(b)** Pupal weight (in mg). **(c)** Adult emergence (%) and **(d)** Adult sex ratio (female:male ratio). Blue: protein-rich diet, Green: standard (balanced) diet, Pink: sugar-rich diet. Black trend line was fitted using the ‘loess’ (non-linear, solid black) and ‘lm’ (linear, dashed black) arguments in R. D1: ~110 larvae/mL diet; D2: ~220 larvae/mL diet; D3: ~450 larvae/mL diet; D4: ~930 larvae/mL diet and D5: ~1800 larvae/mL diet (see Methods for details).

#### Pupal weight

Diet had a statistically significant effect on pupal weight whereby relative to the standard diet, pupae were heavier (lighter) in the protein-rich (sugar-rich) diet (*Diet*: F_2,1033_ = 97.510, p < 0.001, Table 2, Fig 2b). Moreover, density also had a statistically significant effect on pupal weight whereby in general, pupae from higher densities were lighter (*Density*: F_2,1033_ = 1349.130, p < 0.001, Table 2), although the effect of density was non-linear and the decline in pupal weight accelerated more rapidly from densities D3 to D5 (*Density^2^*: F_2,1033_ = 47.655, p < 0.001, Table 2). Importantly, there were statistically significant interaction between diet and the linear (*Diet*Density*: F_2,1033_ = 34.548, p < 0.001) and non-linear effects of density (*Diet*Density^2^*: F_2,1033_ = 9.358, p <0.001). These effects emerged because the linear and non-linear decline in pupal weight with increased density was more pronounced and less quadratic (i.e., closer to linear) in the sugar-rich diet compared with standard and protein-rich diets (Fig 2b).

#### Adult emergence

Diet had a statistically significant effect on adult emergence whereby relative to the standard and protein-rich diets, adult emergence was lower in the sugar-rich diet (*Diet*: F_2,96_ = 42.157, p < 0.001, Table 2, Fig 2c). Moreover, density also had an effect on adult emergence whereby in general, emergence was lower in higher densities (*Density*: F_2,96_ = 152.68, p < 0.001, Table 2), Importantly, there were statistically significant interaction between diet and the linear (*Diet*Density*: F_2,96_ = 23.694, p < 0.001, Table 2) and non-linear effects of density (*Diet*Density^2^*: F_2,96_ = 5.285, p < 0.001). This is because the linear and non-linear decline in adult emergence with increased density was more pronounced and more quadratic (i.e., less linear) in the sugar-rich diet compared with standard and protein-rich diets (Fig 2c).

#### Sex ratio

There were no statistically significant effects of diet, linear and quadratic effects of density, nor their interactions on adult sex ratio (Table 2), whereby the ratio of female:male was similar across densities and diets (Fig 2d).

### Density-by-diet effects on adult traits

#### Body weight

Diet had a statistically significant effect on adult body weight whereby relative to the standard diet, adults were heavier in protein- and sugar-rich diets (*Diet*: F_2,1026_ = 50.087, p < 0.001, Table 2, Fig 3a). Density also had a statistically significant effect on adult body weight whereby adults were in general lighter in higher densities (*Diet*: F_2,1026_ = 1047.71, p < 0.001, Table 2). Sex had statistically significant effect on adult body weight, whereby females were in general heavier than males (*Sex*: F_1,1026_ = 7.569, p = 0.006, Table 2). This effect was also complemented by the statistically significant interaction between density and sex (*Density*Sex*: F_1,1026_ = 5.073, p = 0.002, Table 2), whereby in general the decrease in adult body weight was more pronounced in females than males (Fig 3a). There were also statistically significant interactions of diet and linear (*Diet*Density*: F_2,1026_ = 22.886, p < 0.001, Table 2) and non-linear effect of density (*Diet*Density^2^*: F_2,1026_ = 9.229, p < 0.001, Table 2) whereby the linear decrease in adult body weight with increasing densities was stronger and closer to linear in sugar-rich diet relative to standard and protein-rich diets, with the latter two diets displaying no statistically significant differences (Fig 3a).

**Figure 3.**
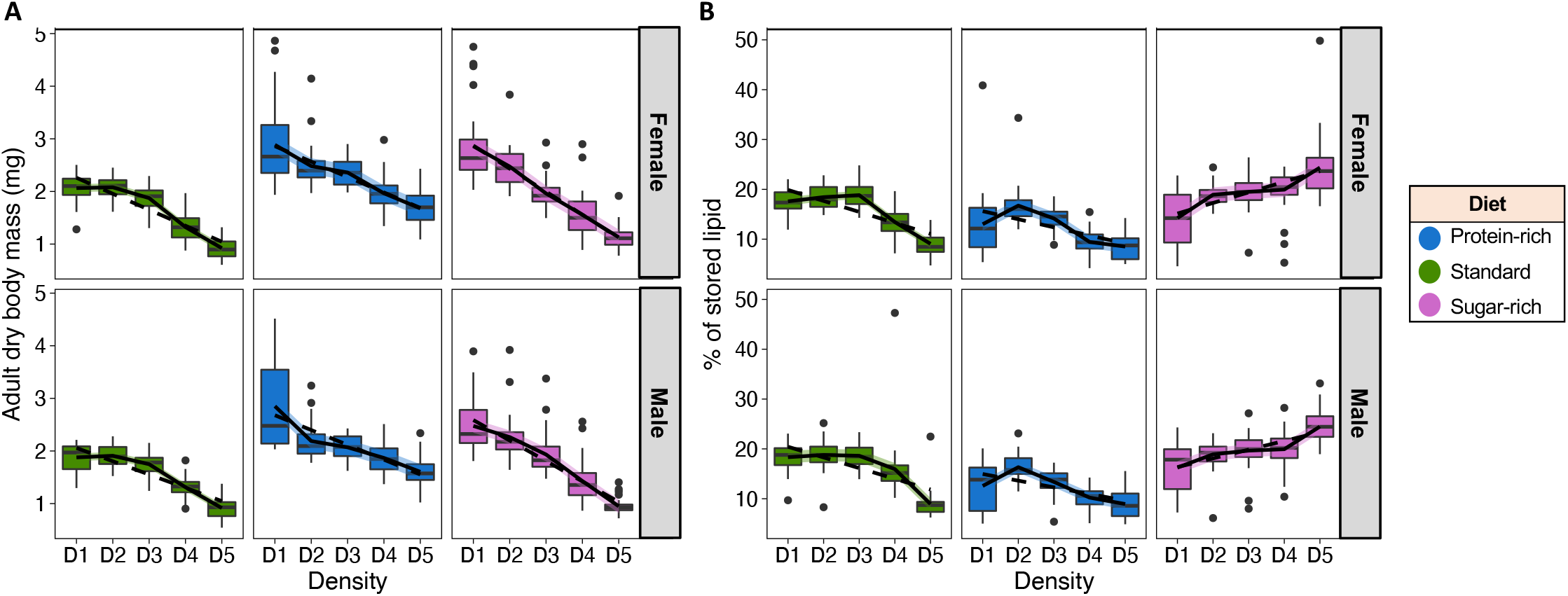
Density-by-diet effects on adult traits. **(a)** Adult dry body mass (mg) and **(b)** Adult lipid reserves (as % of stored lipid relative to body mass). Blue: protein-rich diet, Green: standard (balanced) diet, Pink: sugar-rich diet. Black trend line was fitted using the ‘loess’ (non-linear, solid black) and ‘lm’ (linear, dashed black) arguments in R. D1: ~110 larvae/mL diet; D2: ~220 larvae/mL diet; D3: ~450 larvae/mL diet; D4: ~930 larvae/mL diet and D5: ~1800 larvae/mL diet (see Methods for details).

#### Lipid reserves

Diet had a statistically significant effect on adult lipid reserves whereby relative to the standard diet, adults had higher (lower) lipid reserves in the sugar-rich (protein-rich) diet (*Diet*: F_2,1024_ = 146.640, p < 0.001, Table 2, Fig 3b). Density also had a statistically significant effect on adult lipid reserve whereby in general, adults had lower lipid reserves with increasing densities (but see below; *Density*: F_2,1026_ = 26.023, p < 0.001, Table 2). Sex had no statistically significant effect on lipid reserves, nor had the interactions of sex with density and/or diet (Fig 3b), suggesting that males and females responded similarly to diets and densities. We did find statistically significant interactions between diet and the linear (*Diet*Density*: F_2,1024_ = 115.013, p < 0.001) and non-linear effect of density (*Diet*Density^2^*: F_2,1024_ = 5.869, p < 0.001, Table 2), whereby lipid reserves had a linear *positive* relationship with density in sugar-rich diet, but negative and non-linear relationship with density in standard and protein-rich diets (Fig 3b).

### Density-by-diet effects on adult reproduction

#### Reproductive rates

Diet (*Diet*: F_2, 2057.74_ = 145.36, p < 0.001) and density (*Density*: F_1,2057.74_ = 22.981, p < 0.001; *Density^2^*: F_1,2057.74_ = 8.898, p = 0.002, Table 2, Fig 4a-b) influenced reproductive rates (i.e., daily egg production per female). The main linear effect of day (but not the main non-linear effect of day) was also statistically significant (*Day*: F_1.25.44_= 18.919, p< 0.001; *Day^2^*: F_1, 17.53_ = 3.519, p = 0.077, Table 2). More importantly, there were statistically significant interactions of diet and the linear and quadratic effects of density (*Diet*Density*: F_2, 2057.74_ = 4.744, p = 0.008; *Diet*Density^2^*: F_2, 2057.74_ = 7.904, p < 0.001) as well as diet and the linear and quadratic effects of day on reproductive rate (*Diet*Day*: F_2, 2057.74_ = 68.357, p < 0.001; *Diet*Day^2^*: F_2, 2057.74_ = 50.494, p < 0.001, Table 2, Fig 4a-b). There was also a statistically significant interaction between the linear effect of density and the linear and quadratic effects of day (*Density*Day*: F_1, 2057.74_ = 7.828, p = 0.005, *Density*Day^2^*: F_1, 2057.74_ = 7.768, p = 0.005, Table 2). Overall, these effects emerged from the fact that peak reproductive rates lie at around day 12 and is more evident in low densities in standard diets while in protein- and sugar-rich diets, reproductive rates were earlier in life (~day 1 and day 5, respectively; Fig 4a-b). Groups with individuals form sugar-rich diets had higher reproductive rate at low density relative to groups with individuals from standard diet in low densities (~35 and ~28 daily egg production per female, respectively). Yet, groups with individuals from the standard diet maintained an overall higher reproductive rate throughout the reproductive rate compared with groups with individuals from the sugar-rich diet (Fig 4a). There was also a three-way interaction between diet, linear effect of density and the linear effect of day (*Diet*Density*Day*: F_2, 2057.74_ = 4.317, p = 0.013, Table 2) revealed by contour lines in which non-linear effect of density and day on reproductive rate were pronounced in sugar-rich and standard diets, but became linear in protein-rich diet (i.e., the parallel straight contour lines) (Fig 4a). This means that while increasing density generates a linear decrease in reproductive rate for a given day in protein-rich groups, this relationship is quadratic in groups from sugar-rich and standard diets (Fig 4a-b). Together, the data reveals a complex interaction of diet, density and timing of reproduction shaping groups’ reproductive landscapes.

**Figure 4.**
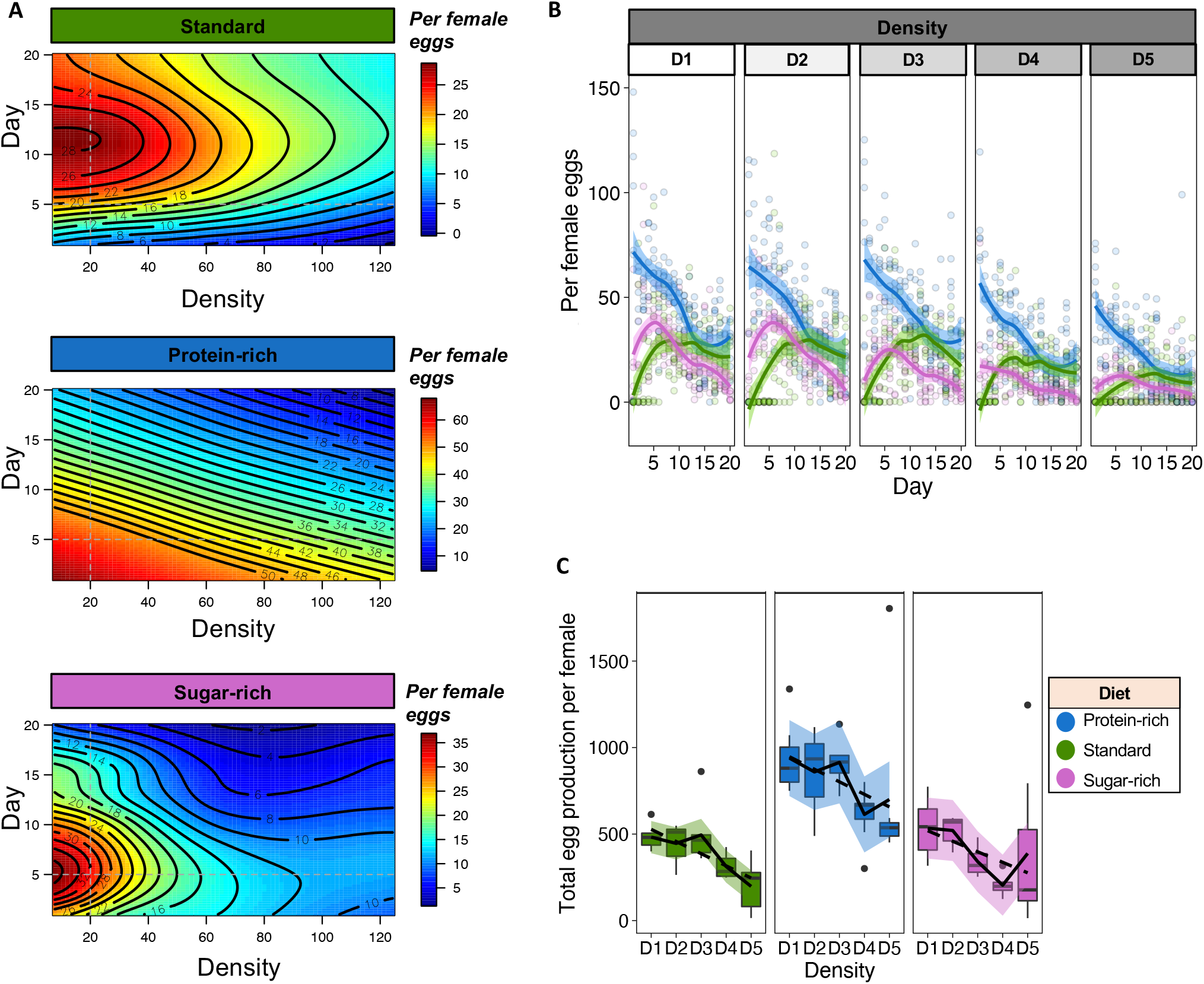
Density-by-diet effects on reproduction. **(a)** Reproductive rate landscape (i.e., egg per female per day). Note that the y-axis presents day, the x-axis represents density, and reproductive rate is the third dimension where high reproductive rates are represented by bright red and low reproductive rates, by dark blue. Note that we used interpolation from the ‘Tps’ function of the ‘fields’ package (see Methods) to generate the surfaces. Dashed grey lines represents changes in reproductive rate for a fixed density (vertical) and day (horizontal), which can be interpreted as in **(b)** non-interpolated data representation of per female daily egg production. Note the peaks in reproductive rate for sugar-rich and standard diets (particularly in low larval densities) which is absent in protein-rich diet, where the relationship is essentially linear. **(c)** Reproductive success (i.e., total egg production per female). In panels (b) and (c): Black trend line was fitted using the ‘loess’ (non-linear, solid black) and ‘lm’ (linear, dashed black) arguments in R. D1: ~110 larvae/mL diet; D2: ~220 larvae/mL diet; D3: ~450 larvae/mL diet; D4: ~930 larvae/mL diet and D5: ~1800 larvae/mL diet (see Methods for details).

#### Reproductive success

Diet had a statistically significant effect on group reproductive success (i.e., total per female egg production) whereby relative to the standard and sugar-rich diets, groups in protein-diet produced overall more eggs (*Diet*: F_2,88.17_ = 22.461, p < 0.001, Table 2). Density also had both linear and non-linear statistically significant effects on group reproductive success whereby groups produced progressively fewer eggs with increasing densities up to the intermediate density D3, after which the negative effect of density on reproductive success was more pronounced (*Density*: F_1,88.05_ = 35.206, p < 0.001; *Density^2^*: F_1,88.202_ = 6.983, p = 0.010, Table 2). There was also a statistically significant interaction between diet and the non-linear effect of density (*Diet*Density^2^*: F_1,88.199_ = 3.177, p = 0.047, Table 2) largely driven by the non-linear decrease in reproductive success with density for groups with individuals from standard diet compared to a relatively more linear decrease in reproductive success with density for groups with individuals from protein-rich and sugar-rich diets (Fig 4c). There were no statistically significant interactions between diet and the linear effect of density.

## Discussion

Conditions experienced early in life modulate individual fitness in adulthood. In holometabolous insects, such as fruit flies, the ecology of the larvae, particularly intraspecific competition and diet, are key factors shaping individual growth and reproduction as adults. In this study, we presented evidence for the interactive effects of larval density and diet in shaping fitness-related traits of the polyphagous fruit fly *B. tryoni*. Our results have overall confirmed our predictions (see Table 1), except for Prediction 4 since our findings show that dietary protein content does not fully rescue the negative effects of high larval density. In fact, in many cases such as for example pupal weight, we found that while a protein rich diet resulted in heavier pupae (Fig 2b), the linear and non-linear effects of density remained similar across diets. This suggests that larval intraspecific competition triggers physiological mechanisms above and beyond protein limitation. One mechanism through which larval density can affect larval growth is through waste. Increasing larval density is known to result in increasing toxicity of the media with compounds such as uric acid and ammonia (Botella et al. 1985). In *D. melanogaster*, high concentrations of ammonia can result in delays in developmental time, increase larval mortality, and larval developmental arrest (Botella et al. 1985). This imposes strong selective pressures on individuals experiencing high larval density conditions which can in turn support polymorphism maintenance (Lewontin 1955; Budnik et al. 1971) particularly in genes related to waste tolerance (Budnik and Brncic 1975). In the long-term, this can assist population to evolve and withstand higher toxicity levels generated by increasing larval densities (Borash et al. 1998). Thus, it is possible that the negative effects of increasing larval densities in our study was derived from the toxicity effects (which are thought to be independent of protein limitation). In our study, we did not quantify toxicity of the media directly, and we do not know whether the genetic makeup of flies in lower and higher densities – and across diets – were different. This remains the main topic for future research in our group.

Our data revealed that protein-rich diets resulted in higher expression of fitness-related traits in high densities (e.g., higher fecundity) (in accordance to Prediction 2) with the exception of the percentage of lipid storage (discussed below). The decrease in fitness trait expression with increasing densities agrees with patterns observed in other insects (Than et al. 2020), invertebrates [e.g., (Zachar and Neiman 2013)] as well as bacteria (Darch et al. 2012) through to vertebrates [e.g., (Svensson et al. 2001)], corroborating the wide taxonomic relationship between population density and fitness. Surprisingly though, we found that while lipid reserve was negatively associated with larval density in protein-rich and standard diet (as expected), lipid reserves increased with larval density in sugar-rich diets. The mechanisms underpinning this effect is unclear and there is no evidence of larval cannibalism [as in *Drosophila* species (Vijendravarma et al. 2013; Kakeya and Takahashi 2020)], but it is possible that protein limitation and high intraspecific competition combined triggered stress responses that resulted in individuals having lower body weight but higher lipid storage. In mice, stress is known to stimulate lipid synthesis but deactivate protein synthesis (Fu et al. 2011)]. Similar stress-related effects could underpin our findings, although further molecular and ecological research are needed to explain the mechanisms.

We found important differences in both the total reproductive success and the reproductive rate curves of populations of flies from varying diets and densities (Fig 4). Overall, populations of flies from protein-rich and/or low densities had relatively higher reproductive success than populations of flies from other diets and/or higher densities (Fig 4c). Interestingly, the shapes of the population reproductive rate curve were markedly different between diets. Populations reared on protein-rich diets reproduced fast and early with the maximum reproductive rate early in lifespan (e.g., before day five) followed by a linear decrease in reproductive rate with age. Conversely, populations reared on sugar-rich and balanced (standard) diets had marked peaks in reproductive rate that were either early in life (e.g., around day five for sugar-rich diets) or mid-life (e.g., around day 10-15 for standard diet) (Fig 4b). The same overall patterns were observed across densities but the effect size decreased with increasing densities, suggesting that density and diet interact to determine group reproductive rate (Fig 4b). Hence, the interaction between diet and density changed the reproductive rate landscape of all populations, resulting in populations with a linear decrease in reproductive rate with age (protein-rich) to populations with early-life and narrow or mid-life and wide peaks in reproductive rate (sugar-rich and standard, respectively) (Fig 4a). What could be the adaptive significance of early reproductive investment for *B. tryoni* populations with high-density individuals and/or from protein-rich diets? Throughout the reproductive season, decaying fruits offer substrates for oviposition and larval development that are progressively richer in protein (e.g., microbial growth) (Silva-Soares et al. 2017). Thus, the protein content of the substrate might function as a phenological cue for the end of the reproductive. For instance, individuals may perceive a short window of reproductive opportunities due to the phenological cue provided by high-protein rotting fruits, which may favour current *vs* future reproductive opportunities; this would explain our findings that *B. tryoni* populations with individuals from protein-rich diets (above and beyond density) display linearly declining reproductive rates (Fig 4b). *B. tryoni* is known to be highly seasonal in its original distribution range (Muthuthantri et al. 2010) opening the possibility for phenological cues to influence *B. tryoni* behaviour. A similar pattern of early life reproduction have been observed in *D. melanogaster* for populations of individuals from high *versus* low larval densities, whereby high-density populations invested in reproduction early in life with a linear decline with reproductive age whereas low-density populations displayed a marked peak in reproductive rate (Morimoto et al. 2017). In insects, larval density is known to act as an ecological cue that shapes adult reproductive physiology and behaviour [e.g., (Johnson et al. 2017)] and changes in ecological factors modulate phenological patterns that can in theory act as cue of the conditions that affect individual reproduction (Abarca and Spahn 2021). It is therefore possible that the high protein diet and/or high larval density (the findings here and in Morimoto et al. 2017) act as cues for faster reproduction, both because of the phenological availability of suitable substrates as well as for the intraspecific competition levels during development. If we are correct, other predictions then emerge with regards to shifts in reproductive rate in response to other ecological factors. For example, we would expect individuals that experienced higher developmental temperatures and/or lower humidity to display a similar reproductive rate curve to those described for high-density individuals and individuals from protein-rich diets. Those factors could act as signals for the end of the optimum reproductive season when the quality of oviposition substrates changes (e.g., higher proportion of rotting fruits) and larval competition in these substrates is intensified. Temperature and rainfall are known to be correlated with the seasonality of *B. tryoni* (Muthuthantri et al. 2010), and developmental temperature is known to accelerate (resting) metabolic rates in *D. melanogaster* (Alton et al. 2020), which could also accelerate reproductive rate. In fact, wild *D. melanogaster* adults sampled in the hottest months of the season displayed higher (maximum) fecundity (Behrman et al. 2015), suggesting that high temperatures during development result in fast-reproducing adults in this species. Future studies that test our hypotheses will help our understanding of the underlying mechanisms resulting in changes in reproductive rates observed in this study and previously, thereby providing insight into anticipatory responses to conditions during development.

In holometabolous insects with little parental care (such as fruit flies), oviposition choices are essential to ensure direct and indirect fitness of current and future generations. The preference-performance hypothesis states that female oviposition choices reflect the performance of the offspring (Levins and MacArthur 1969; Thompson 1988) and many empirical studies corroborate this [see e.g., metanalysis in (Gripenberg et al. 2010)]. It has even been suggested that females can trade optimal foraging patches for better oviposition substrates to the offspring (Lihoreau et al. 2016). Using the preference-performance hypothesis, we can infer that *B. tryoni* females are expected to oviposit preferentially on diets with high-protein content given that protein-rich substrates support faster larval development and greater reproductive success even in high-density conditions relative to other diets. This is assuming that the diet nutritional composition does not change over time. Alternatively, if the composition of the oviposition substrate does change over time, then we can predict females to lay eggs in substrates that will best support offspring growth accounting for this change. In this context, our previous findings suggest that microbial growth significantly improves the quality of sugar-rich diets for larval development (Nguyen et al. 2019). A previous study in *D. melanogaster* has shown that females do oviposit preferentially on sugar-richer diets in the absence of a choice in oviposition substrate (Lihoreau et al. 2016), although *D. melanogaster* larvae develop faster, and with higher reproductive potential, in protein-rich diets compared to carbohydrate-rich diets in laboratory conditions (e.g., with minimal bacterial growth) (Rodrigues et al. 2015). The oviposition behaviour of *B. tryoni* females with regards to the substrate nutritional and bacterial composition is yet unknown and remains a topic for future studies. This will provide insights into the application of the preference-performance hypothesis in this economically important polyphagous fly.

## Conclusions

Holometabolous insects occupy a wide variety of niches, utilising a wide range of substrates for their development. Yet, very little is known about the ecology of larval development in natural habitats, particularly in flies (perhaps with the exception of species with forensic interest). We have recently given the first step towards addressing this gap, by providing the first direct study of *Drosophila melanogaster* larval density in nature (Morimoto and Pietras 2020) but more research is needed, both in the field as well as in controlled laboratory conditions. For instance, the quality of habitats (both across regions and within a region) varies, and so does the quality of the micro-habitat that the larvae experience within the substrate where they develop. The consequences of these differences can be carried over to the adult stage, as well as to the next generations, ultimately contributing to the evolutionary trajectories of groups and populations. In this study, we showed how larval density and larval diet, two important ecological factors in holometabolous insect development, interact to shape life-history trait expression in a polyphagous fruit fly of economic and ecological significance. More broadly, recent studies have demonstrated the role of inter- and trans-generational effects (both in terms of larval diet and density) across some holometabolous insects species [e.g., (Valtonen et al. 2012; Saastamoinen et al. 2013; Woestmann and Saastamoinen 2016; Mbande et al. 2020; Pei et al. 2020)], but a full ecological understanding of the short- and long-term consequences of the quality of larval developmental habitats remains an open field of research. Addressing this gap will be useful to better understand and, ultimately, estimate the relative contributions of each life-stage habitat to individual trade-offs and the overall population dynamics (Stevens et al. 1999, 2000; Monaghan 2008; Spencer et al. 2010).

## Authors’ contributions

JM and FP designed the experiment. AT, BN, IL and HD collected the empirical data. FP and JM supervised the project. JM analysed the data and wrote the draft of the manuscript. All authors contributed to the revision of the manuscript, and approved the final version submitted to the journal.

## Acknowledgements

We thank Prof Owen Lewis and Dr Ana Payo-Payo for helpful comments on the early versions of this manuscript. We thank to MQ-VIED Joint Scholarship for supporting Anh The Than for his study program at Macquarie University. Binh Nguyen was funded by the international Research Training Program (iRTP) scholarship from Macquarie University and the Australian Government. Hue Dinh was funded by Macquarie Research Excellence Scholarship Program scholarship (iMQRES). This research was conducted as part of the SITplus collaborative fruit fly program. Project *Raising Qfly Sterile Insect Technique to World Standard* (HG14033) was funded by the Hort Frontiers Fruit Fly Fund, part of the Hort Frontiers strategic partnership initiative developed by Hort Innovation, with co-investment from Macquarie University and contributions from the Australian Government. The ideas in this manuscript were conceived during the lockdown period of the COVID19 pandemics.

## Data availability

Raw data is available in Dryad: https://doi.org/10.5061/dryad.2280gb5tc and R script to replicate the analysis is available in Zenodo: https://doi.org/10.5281/zenodo.5727007.

